# Temporal regularity shapes thalamocortical responses and deviance sensitivity in the anesthetized rat

**DOI:** 10.64898/2026.11.18.767747

**Authors:** HyunJung An, Fei Peng, Sonja A. Kotz, Jan W.H. Schnupp, Ryszard Auksztulewicz

## Abstract

The brain uses regularities in the sensory environment to anticipate upcoming events, but whether predictions about stimulus identity and timing rely on dissociable neural mechanisms remains unclear. Here, we independently manipulated the temporal structure of acoustic sequences (rhythmic vs. jittered) and stimulus deviance of sequence elements (deviants vs. standards) while recording local field potentials (LFPs) and multiunit activity (AMUA) simultaneously from ventral and dorsal medial geniculate body (MGB) and cortical layers in 9 female anesthetized rats. Temporal structure robustly modulated neural responses throughout the auditory thalamocortical system. LFP responses were stronger for jittered than rhythmic stimulation across all layers of both primary and non-primary auditory cortex, with effects spanning early and later poststimulus intervals. AMUA responses showed a complementary temporal-predictability effect in both dorsal and ventral MGB, with stronger responses to jittered than rhythmic stimuli. In contrast, stimulus deviance effects were strongly dependent on temporal context and recording site. Deviant–standard differences were enhanced under jittered stimulation in cortex and in early MGB responses, whereas later MGB responses showed a context-dependent reversal. Finally, AMUA-based Granger analyses revealed sustained jitter-specific directed influences from both dorsal and ventral MGB toward multiple cortical layers during the poststimulus period. Together, these findings demonstrate that temporal regularity robustly modulates auditory thalamocortical responses and is associated with changes in directed interactions between thalamus and cortex. Responses to stimulus deviance, in turn, depend on temporal context and recording site. These findings reveal distinct but interacting neural signatures of temporal regularity and stimulus deviance in the auditory thalamocortical system.

## INTRODUCTION

How the brain uses regularities in the sensory environment to anticipate incoming events and update expectations when they are violated is a central question in neuroscience. In addition, this predictive capacity is commonly studied by examining neural responses to unexpected sounds. A standard approach is the oddball paradigm, in which rare deviants are contrasted with frequent standards, eliciting mismatch responses (MMRs) (Näätänen et al. 2007; Garrido, Kilner, Kiebel, et al. 2009; Garrido, Kilner, Stephan, et al. 2009; Heilbron and Chait 2018). Within predictive-processing accounts, such responses are often interpreted as reflecting prediction-error signalling when incoming sensory input deviates from an internal model of the environment (Garrido, Kilner, Kiebel, et al. 2009; Garrido, Kilner, Stephan, et al. 2009; Friston 2005; Wacongne et al. 2012). However, standard–deviant contrasts can also include contributions from stimulus-specific adaptation, repetition history, and other sensory-context effects. An important question is therefore not only where mismatch-related signals arise, but whether distinct forms of sensory regularity engage dissociable neural mechanisms.

Regularities in auditory sequences can concern both the identity of an event (what sound is expected) and its timing (when it is expected). Although these dimensions often covary in natural sensory environments (Ten Oever and Martin 2024), they can be distinguished conceptually and experimentally (Nobre and van Ede 2017; Heilbron and Chait 2018). Evidence from human electrophysiology, neuroimaging, and intracranial recordings suggests that ‘what’ and ‘when’ predictability can influence auditory processing through partly distinct mechanisms. In particular, ‘what’ and ‘when’ predictions have been associated with separable forms of short-term plasticity and gain control in auditory and sensorimotor systems (Auksztulewicz et al. 2018). However, these findings do not establish whether identity-related deviance and temporal regularity are processed by distinct components of the auditory thalamocortical system. Animal electrophysiology provides an opportunity to examine how sensitivity to these two dimensions is expressed across thalamic subdivisions and cortical layers.

A potential substrate for differentiated processing of content and temporal structure is provided by the organization of auditory thalamocortical circuitry (Musacchia et al. 2014). The auditory system comprises partially segregated lemniscal and non-lemniscal pathways that differ in connectivity, physiology, and functional organization (Jones 1998, 2001; Jones 2003; de Venecia et al. 1995; Winer et al. 2005; Lee and Winer 2008; Lomber and Malhotra 2008). The lemniscal auditory pathway enters the forebrain primarily through the ventral division of the medial geniculate body (MGBv), which receives ascending input from the central nucleus of the inferior colliculus and projects predominantly to granular layers of primary auditory cortex. MGBv neurons show relatively narrow frequency tuning and preserve precise tonotopic organization, properties well suited to representing acoustic features such as spectral content and sound identity (Calford 1983; McMullen and de Venecia 1993; de Venecia et al. 1995; Pandya et al. 1994; Hashikawa et al. 1995; Kimura et al. 2003; Anderson et al. 2007). By contrast, non-primary auditory thalamic nuclei receive broader and more heterogeneous inputs, tend to show less sharply tuned frequency responses, and project to distinct, partly overlapping cortical fields and laminae (Jones 2001; Rauschecker and Tian 2000). These anatomical distinctions provide a plausible basis for differentiated processing of auditory identity and temporal structure, but they do not establish a strict segregation of content- and timing-related predictions.

However, functional distinctions between ‘what’ and ‘when’ processing do not necessarily imply a corresponding division of predictive computations across lemniscal and non-lemniscal pathways. Stimulus-specific adaptation, whereby responses to repeated sounds are attenuated relative to responses to rare sounds, has been demonstrated in both lemniscal and non-lemniscal divisions of the auditory thalamus, with particularly strong adaptation in non-lemniscal regions (Antunes et al. 2010). More broadly, recordings along the rodent auditory pathway have revealed deviance-related responses in the inferior colliculus, MGB, and auditory cortex, with evidence that repetition suppression and prediction-error responses vary across processing levels and pathway subdivisions (Parras et al. 2017). These findings suggest that auditory deviance sensitivity emerges at multiple levels of the auditory system, while leaving open how temporal regularity modulates these responses across thalamic subdivisions and cortical layers.

Temporal regularity can also shape auditory processing through mechanisms that do not require an explicit behavioral task. In behaving rats, temporal expectations have been shown to modulate sound-evoked responses in the primary auditory cortex and to improve perceptual performance (Jaramillo and Zador 2011). In anesthetized rats, cortical oscillatory activity is sensitive to rhythmic sequence structure, suggesting that the auditory cortex can track temporal regularities even in the absence of task-related attention (Noda et al. 2017). These findings complement accounts in which beat-based entrainment and interval-based timing support temporal expectations through partly distinct mechanisms (Breska and Deouell 2017). Rhythmic processing has also been linked to interactions between thalamocortical inputs and cortical oscillatory networks, potentially modulating cortical excitability at expected moments in time (Schroeder and Lakatos 2009; Lakatos et al. 2007, 2013; Musacchia et al. 2014). Importantly, sensitivity to auditory deviance and temporal regularity has also been demonstrated in anesthetized rats (Parras et al. 2017; Noda et al. 2017; Cappotto et al. 2023), investigating how these regularities shape neural responses across the auditory thalamocortical system. However, it remains unclear how temporal regularity affects responses to stimulus deviance across thalamic subdivisions and cortical layers, and whether these effects are accompanied by changes in directed thalamocortical statistical interactions.

The present study investigated how sound identity and temporal regularity shape neural processing across the auditory thalamocortical system. We independently manipulated identity-related regularity by comparing responses to standard and deviant sounds, and temporal regularity by presenting sounds in rhythmic or temporally jittered sequences. We considered three main predictions. First, if temporal regularity promotes response suppression, rhythmic sequences should elicit smaller neural responses than jittered sequences. We tested whether this effect was expressed across cortical fields and layers and in both dorsal and ventral medial geniculate body (MGB), or whether it differed systematically across these anatomical compartments. Second, if identity-related deviance processing depends on temporal context, the difference between deviant and standard responses should vary between rhythmic and jittered sequences, even when the immediately preceding interstimulus interval is matched. Conversely, a deviant–standard difference that remains similar across temporal conditions would be more consistent with a context-independent identity effect. Third, we asked whether the distribution of identity- and timing-related effects would support a simple anatomical segregation of these processes or instead reveal context-dependent interactions across the thalamocortical system. Preferential localization of temporal-context effects and identity-related effects to different anatomical compartments would be consistent with functional differentiation, whereas significant interactions between stimulus identity and temporal condition would indicate that deviance-related responses depend on temporal context. We further examined whether temporal condition and stimulus identity were associated with changes in directed statistical interactions between thalamic and cortical recording sites. To address these questions, we simultaneously recorded local field potentials (LFPs) and multiunit activity (AMUA) from dorsal and ventral MGB and multiple layers of primary and non-primary auditory cortex in anesthetized rats, and used Granger causality to assess condition-dependent directed statistical interactions. This approach allowed us to characterize both the anatomical distribution of temporal-context modulation and the extent to which identity-related responses depended on temporal regularity.

## MATERIALS AND METHODS

All experimental procedures were approved by the Committee on the Use and Care of Animals at City University of Hong Kong and under license by the Department of Health of Hong Kong [Ref. No. (17–72) in DH/SHS/8/2/5 Pt.1].

### Subjects and Surgical procedures

Nine female adult Wistar rats (8–20 weeks of age, mean = 10 weeks; body weight 200–310 g, mean = 238 g) were obtained from the Chinese University of Hong Kong for this study. Only female rats were used to maintain consistency with previously published experimental protocols using comparable anesthetized-rat preparations (An et al. 2021). Prior to surgery, all animals were assessed for normal auditory function by recording auditory brainstem responses (ABRs). ABRs were elicited by 500 μs white-noise clicks delivered at 23 Hz through hollow ear bars equipped with custom-built headphone drivers (AS02204MR-N50-R, PUI Audio, Dayton, USA). For each intensity level (30–80 dB SPL, 5 dB steps), 400 clicks were presented. Neural activity was recorded with stainless steel needle electrodes positioned at the mastoids, nose, and back, and ABR waveforms were derived from averaged scalp potentials between the mastoid and the vertex (Rosskothen-Kuhl et al. 2021). Animals with ABR thresholds of 30 dB SPL or lower were included as normal-hearing subjects.

At the beginning of each recording experiment, anesthesia was induced via intraperitoneal (i.p.) injection of ketamine (80 mg/kg) and xylazine (12 mg/kg). In addition, all rats received an i.p. injection of the anti-inflammatory dexamethasone (0.2 mg/kg) prior to surgery. Urethane (0.75 g/kg, i.p.) was administered ∼1 h after induction with ketamine and xylazine, and additional doses (0.1–0.2 ml, i.p.) were given as needed to maintain anesthesia depth. Ketamine + xylazine was selected for induction as it provides rapid onset of deep anesthesia, while urethane was used for recordings due to its minimal impact on neural activity (Malmierca et al. 2019).

Thus, all neural recordings were performed under urethane anesthesia following surgical preparation under ketamine/xylazine anesthesia. Anesthetic depth was continuously monitored by assessing the toe-pinch withdrawal reflex. Body temperature was measured rectally and maintained at 36 ± 1 °C using a thermostatically controlled heating pad (RWD Life Sciences, Shenzhen, China). During surgery and recording, the rats were placed in a sound-attenuated chamber and secured in a stereotaxic frame. The hollow ear bars were used both to stabilize the head for craniotomy and to deliver auditory stimuli. Craniotomies were performed at two sites, targeting the MGB and the AC, respectively.

For simultaneous recordings from the auditory cortex (AC) and medial geniculate body (MGB), separate craniotomies were performed over the left hemisphere to provide access to the cortical and thalamic recording sites. For MGB recordings, a craniotomy was performed at approximately 5.5 mm caudal and 3.6 mm lateral to bregma, based on stereotaxic coordinates from the rat brain atlas (Paxinos and Watson 2007). A single-shank, 32-channel silicon probe with 50-μm inter-electrode spacing (ATLAS Neuroengineering, E32-50-S1-L6) was advanced dorsoventrally to approximately 6 mm ventral to bregma to reach the MGB. The probe was positioned to record neural activity from the putative dorsal and ventral subdivisions of the MGB. As probe tracks were not histologically verified, subdivision assignments were based on probe depth and physiological response characteristics (see Channel mapping). The rostrocaudal extent of the targeted MGB region was referenced using coronal sections spanning approximately −5.2 to −5.8 mm along the AP axis, with the primary recording plane centered at AP −5.5 mm. For AC recordings, a 5 × 4 mm craniotomy was created to expose the left auditory cortex, extending approximately from 2.5 to 7.5 mm posterior to bregma, with the medial edge located approximately 2.5 mm lateral to the midline (Polley et al. 2007). Neural activity was recorded using a 64-channel silicon probe consisting of four parallel shanks, with 16 recording sites on each shank and 100-μm inter-site spacing (ATLAS Neuroengineering, E64-100-S4-L6-600). The probe was inserted approximately perpendicular to the cortical surface, allowing the recording sites to span the cortical depth from superficial to deep layers and thereby preserve laminar information across the cortical column.

Recording sites were subsequently classified according to their anatomical position and functional response properties. Cortical recording sites were assigned to three laminar compartments according to recording depth: superficial layers (L2/3), middle (thalamorecipient) layer (L4), and deep layers (L5/6). The middle compartment was defined physiologically by the earliest CSD sink and, because thalamocortical afferents in auditory cortex also reach lower layer 3, need not coincide exactly with cytoarchitectonic layer 4 (Szymanski et al. 2009; Winer et al. 2005). MGB recording sites were categorized as belonging to either the dorsal MGB (dMGB) or ventral MGB (vMGB) according to their dorsoventral position along the probe trajectory and correspondence with stereotaxic atlas coordinates. Functional characterization of the recording sites was performed using auditory response measurements obtained prior to the main experimental sequences, as described in the following section. As electrode placements were not verified histologically, compartment assignments were based on stereotaxic coordinates, probe depth, and physiological response properties. Together, this recording configuration enabled simultaneous assessment of cortical and thalamic neural activity while preserving laminar organization within the AC and dorsoventral subdivision of the MGB (Fig. 1)

**Figure 1.**
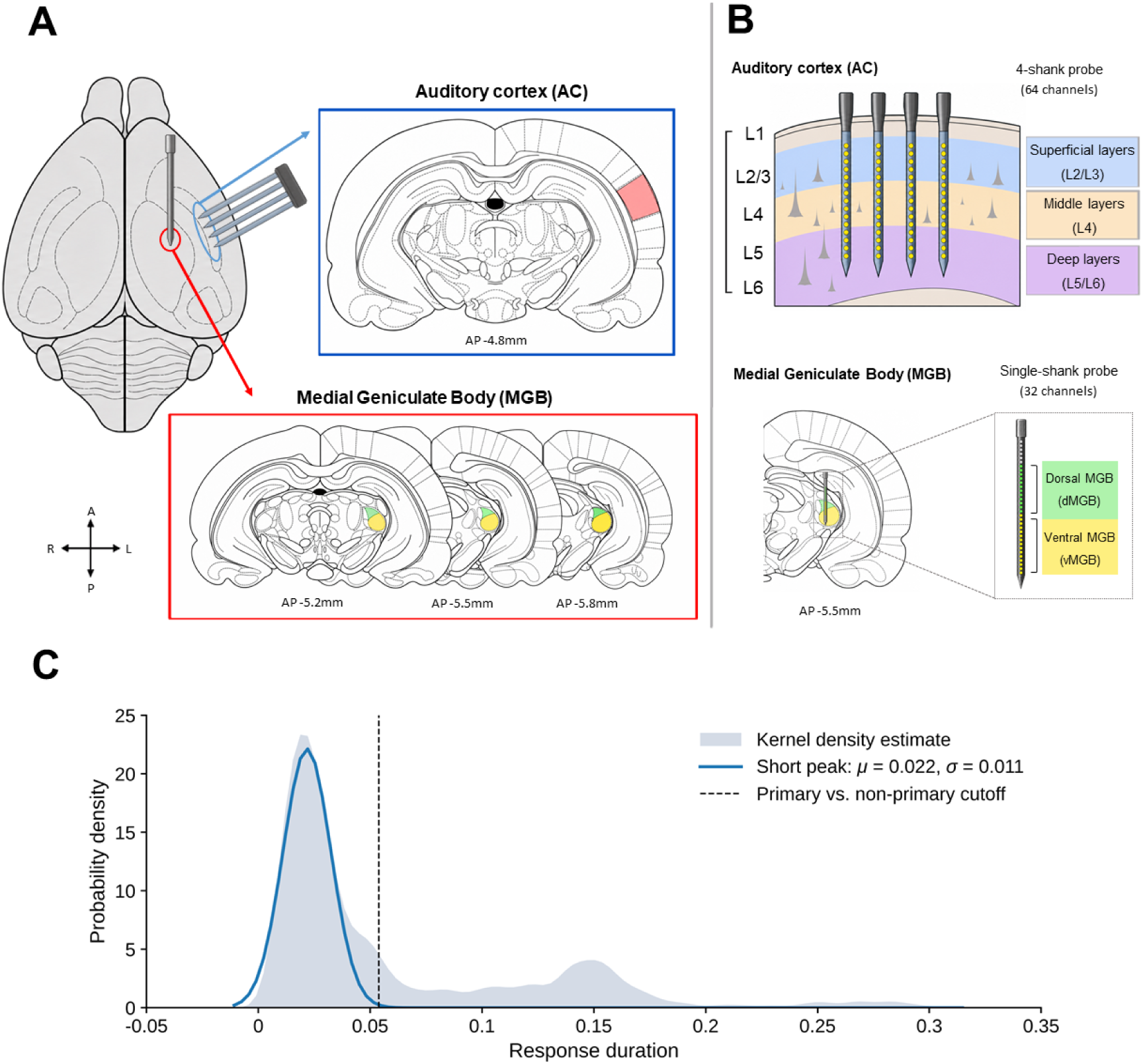
Schematic drawings illustrating simultaneous recording locations and classification of recording sites in the auditory cortex and medial geniculate body. (A) Schematic of the simultaneous recording configuration in the left auditory cortex (AC) and medial geniculate body (MGB). Neural activity in the AC was recorded using a 64-channel, four-shank silicon probe inserted approximately perpendicular to the cortical surface to sample activity across the cortical depth. The enlarged coronal outline (anteroposterior (AP) −4.8 mm; adapted from (Paxinos and Watson 2007) shows the approximate cortical recording location. MGB activity was recorded using a 32-channel single-shank silicon probe inserted dorsoventrally at approximately 5.5 mm posterior and 3.6 mm lateral to bregma. Coronal outlines at AP −5.2, −5.5, and −5.8 mm, redrawn from the rat brain atlas(Paxinos and Watson 2007), indicate the rostrocaudal extent and approximate anatomical location of the targeted MGB region. (B) Schematic of the classification of recording sites used for subsequent analyses. Cortical recording sites sampled by the four-shank probe were assigned to three laminar compartments according to cortical depth: superficial layers (L2/3), middle (thalamorecipient) layer (L4), and deep layers (L5/6). MGB recording sites sampled by the single-shank probe were categorized into dorsal MGB (dMGB) and ventral MGB (vMGB) according to their dorsoventral position along the recording trajectory and corresponding stereotaxic coordinates. Recording sites were assigned to compartments based on stereotaxic coordinates and a functional characterization of the recording sites using responses to white-noise stimulation and frequency response area mapping (see the Auditory paradigm section and Channel mapping sections). (C) Distribution of channel-level response durations used to define the A1/non-A1 classification threshold. The dashed line indicates the cutoff (53.8 ms), defined as the mean + 3 SD of the short-duration Gaussian component.

### Auditory paradigm

To check that channels contained signals from neuronal populations sensitive to acoustic frequency, frequency response areas (FRAs) were measured. For FRA mapping, pure tones (duration 100 ms with 5 ms rise/fall ramps) spanning 0.5–22.627 kHz were presented at 23 logarithmically spaced frequencies in quarter-octave steps. Tones were presented at sound levels ranging from 30 to 80 dB SPL in 10-dB increments, allowing frequency-dependent response profiles to be characterized across recording sites. White-noise (WN) stimulation (100 ms with 5 ms rise/fall ramps) at 80 dB SPL was also used as a broadband search stimulus to verify the presence of sound-evoked neural activity across recording channels. WN and FRA measurements were obtained for every recording electrode position sampled. Auditory stimuli were digitally generated at a sampling rate of 48,828.125 Hz and presented diotically through headphones. Electrophysiological signals were acquired at a sampling rate of 24,414.0625 Hz using a Tucker-Davis Technologies (TDT) System 3 recording setup. Neural responses to WN and FRA stimuli were subsequently characterized using analog multiunit activity (AMUA) and current source density (CSD) measures. For functional channel classification, AMUA responses to all pure tones presented at 70 dB SPL were pooled across frequencies to obtain one frequency-pooled response time course (PSTH) per channel. This PSTH and the WN evoked PSTH were used to assess responsiveness, response onset and response duration; preprocessing and classification criteria are described in the Channel mapping section below.

In the main experimental sequences (Fig. 2), consonant-vowel (CV) syllable stimuli were taken from the pitch condition of our previous study (An et al. 2021). Briefly, two CV syllables, /da/ and /ba/, were selected from a corpus of natural speech syllables (Ives et al. 2005), digitally edited using the STRAIGHT software toolbox (Kawahara 2006) for Matlab (MathWorks Inc., Natick, USA), and blended at a ratio of 0.5:0.5 by mixing their spectrograms. From this syllable, 75 ms stimulus tokens were generated that differed only in fundamental frequency (F0). Five F0 levels were used, centred on 1 kHz and separated by quarter-octave steps (0.7071, 0.8409, 1, 1.1892, and 1.4142 kHz). As harmonic sounds, these syllables contained energy at integer multiples of F0, extending well above the F0 range. The animals had no prior exposure to these stimuli before the experiment. Stimuli were generated at a sampling rate of 44.1 kHz and presented diotically at 65 dB SPL.

**Figure 2.**
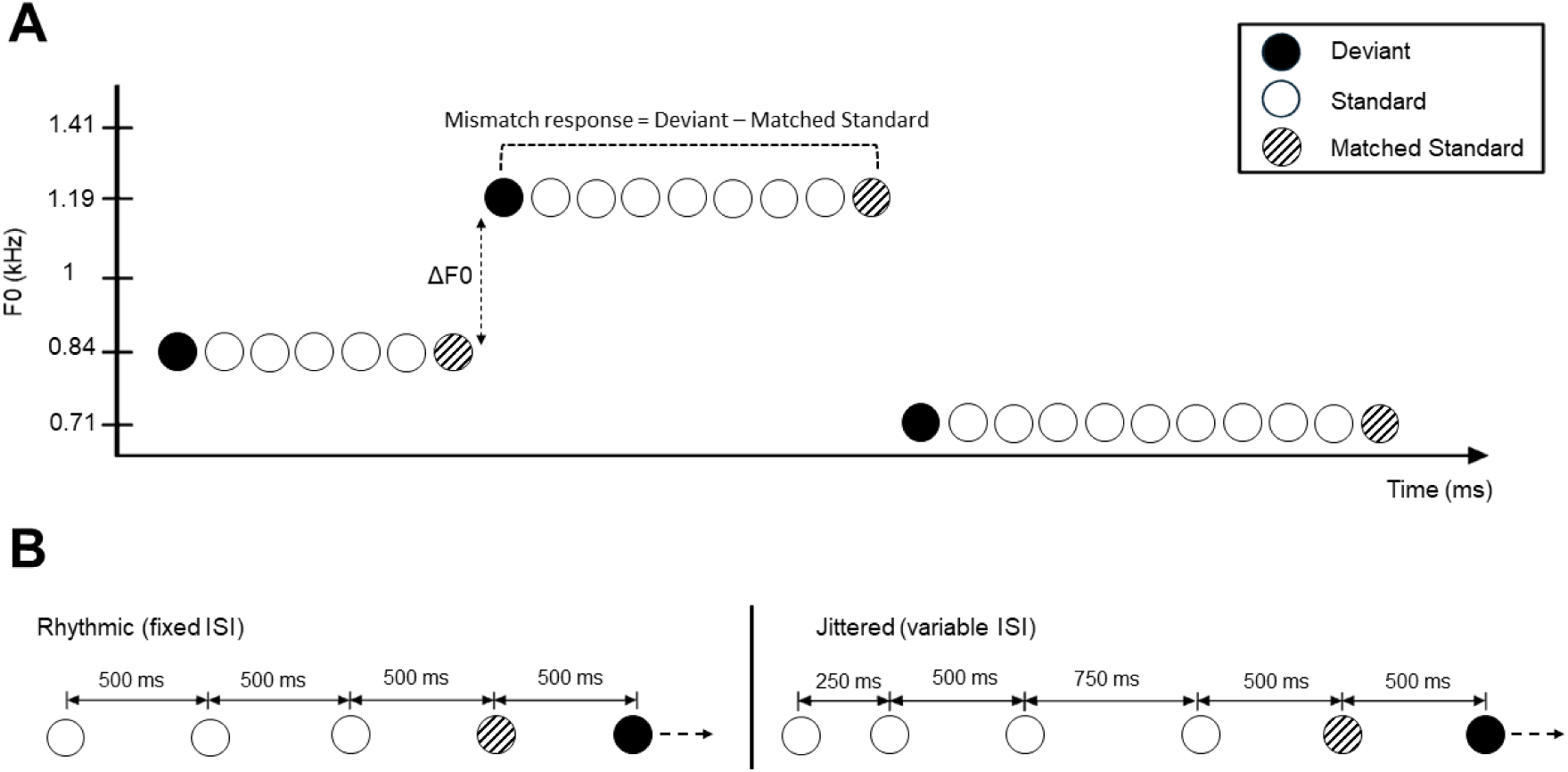
Roving oddball paradigm and temporal conditions. **(A)** Schematic illustration of the roving oddball paradigm. Stimuli consisted of synthetic consonant–vowel syllables (75 ms) presented at five fundamental frequencies (F0: 0.71, 0.84, 1.00, 1.19, and 1.41 kHz; quarter-octave steps) (An et al. 2021) in trains of 7, 9, 11, or 13 stimuli. The first stimulus following a change in F0 was defined as the deviant (filled circles), whereas subsequent repetitions at the same F0 were defined as standards (open circles). Hatched circles represent matched standards selected near the end of each train, each preceded by a 500-ms interstimulus interval (ISI), for comparison with the corresponding deviant. All five F0 values served as both standards and deviants across the experiment. **(B)** Schematic illustration of the two temporal conditions. In the rhythmic condition, stimuli were presented at a fixed ISI of 500 ms. In the jittered condition, ISIs were randomly set to either 250 or 750 ms, except for the intervals immediately preceding each deviant and matched standard, which were fixed at 500 ms. Thus, all stimuli included in the deviant–standard comparisons were preceded by the same 500 ms ISI in both temporal conditions.

A roving oddball paradigm was used to investigate auditory deviance detection. In this paradigm, CV stimuli were presented in continuous trains (Fig. 2A). The first stimulus of each train constitutes an auditory deviant, which becomes a standard after a few repetitions within the train. After 7, 9, 11 or 13 presentations, a new stimulus with a different F0 is chosen, and its first occurrence thus constitutes a deviant relative to the previous train. This stimulus then starts a new train, during which it becomes the new standard. Across the experiment, the five F0s were balanced such that each served equally often as the standard and deviant. In the analysis, responses to the first stimulus of each train (deviant) and a matched standard from the same train (defined below) were compared. This ensures that deviant and standard responses have precisely the same stimulus parameters and number of trials.

To manipulate temporal regularity, roving oddball sequences with identical stimulus structures were presented under two temporal conditions: rhythmic and jittered (Fig. 2B). In the rhythmic condition, stimuli were presented at a fixed inter-stimulus interval (ISI) of 500 ms, producing an isochronous sequence. In the jittered condition, ISIs were randomly set to either 250 or 750 ms, except for the intervals immediately preceding each deviant and one matched standard per train, which were fixed at 500 ms. This manipulation maintained a mean ISI of approximately 500 ms in both conditions while disrupting the regular temporal structure. Importantly, the immediately preceding ISI was identical (500 ms) for all stimuli included in the deviant– standard comparisons. Note, however, that the jittered condition reduced rather than eliminated temporal predictability, because the discrete set of possible ISIs constrained when the next stimulus could occur (e.g., if no stimulus had occurred 250 ms after the previous one, it would occur at 500 or 750 ms).

Rhythmic and jittered conditions were presented in separate blocks, with each condition repeated twice, in a random order that differed across animals. Each block contained 1,600 stimuli organized into 160 trains of 7, 9, 11, or 13 stimuli, yielding 160 deviants and 160 matched standards. The matched standard was the standard preceded by a 500-ms ISI near the end of each train: the last standard in the rhythmic condition, and the last or second-to-last standard in the jittered condition, depending on the train. Thus, each temporal condition comprised 3,200 stimuli in total.

### Channel mapping

Before the main analyses, analog multiunit activity (AMUA) and current source density (CSD) analyses were performed to assign recording sites to functional subdivisions of the auditory cortex (AC) and medial geniculate body (MGB) respectively, according to physiological markers.

For each channel, AMUA was extracted from the broadband signal by band-pass filtering between 500 and 3000 Hz (third-order two-pass Butterworth filter), rectification and downsampling to 500 Hz. The AMUA signal was then epoched relative to stimulus onset and averaged across trials to obtain response time courses (peri-stimulus time histograms, PSTHs). Two PSTHs were computed per channel: one for WN bursts, and one for pure tones presented at 70 dB SPL, pooled across all tested frequencies (“frequency-pooled tone PSTH”). Frequency pooling was used to characterize the presence and temporal profile of tone-evoked responses rather than frequency tuning. PSTHs were baseline-corrected and denoised using principal component analysis (PCA), retaining eight principal components. For each PSTH, we quantified the peak response amplitude, baseline mean and standard deviation (SD), peak-to-baseline SD ratio, and response onset, offset, and duration. Response onset and offset were determined from the first and last threshold crossings, respectively, using a threshold corresponding to 50% of the peak response above baseline. Response duration was calculated as the interval between response onset and offset.

AC electrode shanks were functionally classified as penetrating A1 or non-A1 cortex based on a combination of response amplitude and response duration, reflecting differences in stimulus responsiveness and temporal response properties across auditory cortical fields (Bizley et al. 2005; Kajikawa et al. 2005). Classification was performed at the shank level, assuming that each shank sampled a single cortical field. Channels whose WN AMUA responses did not exceed 3 SD above baseline were excluded. For each shank, we calculated the proportion of WN-responsive channels whose frequency-pooled tone PSTHs did not exceed 3 SD above baseline. Shanks in which more than 50% of WN-responsive channels failed to meet this tone-response criterion were classified as non-A1. For the remaining shanks, classification was based on response duration. For each eligible channel, response duration was averaged across the WN PSTH and the frequency-pooled tone PSTH, and these channel-level averages were pooled across eligible shanks to characterize the distribution of response durations. The distribution was estimated using kernel density estimation and fitted with a Gaussian mixture model (Fig. 1C). The duration threshold was defined as the mean plus 3 SD of the short-duration component, yielding a cutoff of 53.8 ms. For each shank not already classified as non-A1, response duration was averaged across eligible channels and both stimulus conditions. Shanks with mean response durations below 53.8 ms were classified as A1, whereas those with durations above this threshold were classified as non-A1.

Laminar organization within AC was assessed using current source density (CSD) analysis of local field potentials (LFPs). CSD estimates the spatial distribution of transmembrane current sources and sinks from the second spatial derivative of extracellular potentials recorded at different cortical depths. In AC, early stimulus-evoked CSD sinks in the middle (thalamorecipient) layers are commonly interpreted as reflecting thalamocortical input, providing a physiological landmark for estimating laminar position (Happel et al. 2010; Szymanski et al. 2011). LFPs recorded during WN and FRA stimulation were low-pass filtered at 100 Hz and high-pass filtered at 1 Hz using third-order Butterworth filters, baseline-corrected by subtracting the mean voltage during the first 100 ms, and downsampled to 500 Hz. To reduce noise, denoising source separation (DSS) was applied separately to the WN and FRA data (de Cheveigné and Simon 2008; de Cheveigné 2020), and the signals reconstructed from the first eight DSS components were used for subsequent analyses. Responses were averaged across repetitions; for FRA stimulation, responses were additionally averaged across tested frequencies at 70 dB SPL. CSD profiles were calculated separately for WN and FRA responses by taking the second spatial difference of the LFP signals along each shank. For each channel, CSD response onset was defined as the earliest poststimulus time at which the response crossed a threshold equal to half the difference between its peak amplitude and baseline. Onset times obtained from WN and FRA responses were averaged for each channel. Among channels classified as good according to the preceding AMUA-based quality criteria (WN response > 3 SD relative to baseline), the channel with the earliest CSD response was designated the earliest responder. When multiple channels shared the earliest CSD onset, AMUA response onset was used to select the earliest responder; if a tie remained, the candidate channel closest to the middle of the shank was selected. This procedure identified a unique earliest-responder channel for all but two shanks, for which the middle channel was selected. The earliest-responder channel and its two immediate neighbours were designated as the middle (thalamorecipient) layer, corresponding to an operational thickness of approximately 200 μm, consistent with anatomical estimates of granular-layer thickness in rat auditory cortex (Tao et al. 2017). Channels superficial to this region were classified as superficial, and channels deeper than this region were classified as deep. The use of CSD and response onset to identify the middle (thalamorecipient) region follows established approaches for functionally defining cortical laminae from the earliest evoked sink (Ferro et al. 2021).

MGB recording sites were processed using the same AMUA pipeline as AC sites. The functional subdivision of the MGB into ventral and dorsal regions was guided by converging differences in response amplitude, response-onset latency, and electrode depth. In particular, responses to WN stimulation showed longer onset latencies in putative dorsal MGB than in putative ventral MGB, consistent with previous physiological studies reporting distinct response properties in these regions (Anderson and Linden 2011). To operationalize this subdivision, the electrode exhibiting the largest peak response was used as a functional reference point. Based on the approximate 600-μm dorsoventral extent of the rat ventral MGB, sites within ±300 μm of this reference electrode were classified as ventral MGB (vMGB), whereas sites located 300–900 μm dorsal to the reference were classified as dorsal MGB. Thus, site assignment combined physiological response characteristics with electrode depth to provide an operational estimate of the MGB subdivisions.

### Analysis of deviants and standards in rhythmic and jittered sequences

For the analysis of neural responses to deviants and standards presented in rhythmic and jittered sequences, LFP signals were obtained from the continuous recordings using the corresponding low-frequency signal component and were downsampled to 500 Hz. AMUA was extracted by band-pass filtering the signals between 500 and 3000 Hz using second-order Butterworth filters applied in both forward and reverse directions, followed by computation of the root-mean-square (RMS) envelope using a 30-sample window. Both LFP and AMUA signals were epoched from −100 to 400 ms relative to stimulus onset and baseline corrected by subtracting the mean activity during the 100 ms preceding stimulus onset. For each recording site and temporal condition, the analysis was restricted to events preceded by a 500-ms ISI to match the immediately preceding interval across rhythmic and jittered sequences.

To reduce noise and non-neural contributions, LFP and AMUA responses were separately denoised using DSS. DSS was performed separately for the stimulus-locked standard and deviant responses and for the complete set of single-trial epochs. For each dataset, covariance matrices across all channels were computed from the single-trial data and from the trial-averaged responses, and a DSS transformation was derived to identify spatial components with enhanced reproducibility across trials. The first eight DSS components, which captured the most reproducible stimulus-related activity, were retained and projected back into channel space using the inverse transformation. The resulting denoised data were used for subsequent analyses of standard and deviant responses in the rhythmic and jittered conditions.

Statistical analyses were performed separately on LFP and AMUA response time series. Baseline activity was pooled across standard and deviant trials from both temporal conditions, and the standard deviation was calculated across baseline time points of the trial-averaged signal for each channel. The resulting normalization factor was then used to scale the time series of each trial within the corresponding channel. This procedure expressed response amplitudes relative to baseline variability (averaged across conditions) and facilitated comparisons across channels with different signal scales. AMUA time series were additionally smoothed using a five-point moving window. Statistical analyses were conducted on time points between −50 and 250 ms relative to stimulus onset. The pre-stimulus interval from −50 to 0 ms was included to assess activity before stimulus onset, while the post-stimulus interval ended at 250 ms to avoid potential contamination from the next stimulus, given that the shortest inter-stimulus interval in the jittered condition was 250 ms.

We tested whether neural responses differed between standard and deviant sounds, whether responses differed between rhythmic and jittered sequences, and whether the deviant–standard difference depended on temporal condition. A deviant–standard difference is consistent with sensitivity to sound-identity regularity, whereas an interaction between stimulus type and temporal condition indicates that this difference varies with temporal context. Because the immediately preceding ISI was matched across conditions, such an interaction cannot be attributed simply to a difference in that interval. We also tested whether these effects varied across recording regions.

To characterize the time course of these effects and test whether they differed across the auditory thalamocortical system, linear mixed-effects models were fitted independently at each peri-stimulus time point. The dependent variable was the normalized neural response, with fixed effects of stimulus type (deviant vs. standard), temporal condition (rhythmic vs. jittered), recording region, and their interactions. For cortical recordings, recording categories were defined by the combination of auditory field (primary vs. non-primary) and cortical layer (superficial, middle (thalamorecipient), or deep), yielding six cortical categories; ventral and dorsal MGB constituted two additional categories. The stimulus-type effect quantified the deviant–standard response difference, and the temporal-condition effect quantified the rhythmic–jittered difference. The stimulus-type × temporal-condition interaction tested whether the deviant–standard difference varied with temporal condition. Interactions involving recording regions tested whether these effects differed across the thalamocortical recording categories. Recording penetration was included as a random effect to account for correlations among measurements obtained from the same penetration, rather than treating those measurements as independent. A random intercept accounted for baseline differences between penetrations, while random slopes for the stimulus-type × temporal-condition interaction allowed this interaction to vary across penetrations.

To determine which recording regions contributed to the effects identified by the omnibus analysis, we next tested whether stimulus type, temporal condition, and their interaction were associated with responses within each of the eight recording categories. These region-specific models assessed whether the deviant–standard difference, the rhythmic–jittered difference, and the dependence of the former on temporal condition were present within each category. Separate linear mixed-effects models were fitted for each category at each time point. Stimulus type, temporal condition, and their interaction were included as fixed effects, and recording penetration was included as a random effect, with a random intercept and random slopes for the stimulus-type × temporal-condition interaction. This approach accounted for non-independence among measurements obtained from the same penetration while allowing the effects of interest to be assessed separately within each recording category.

For each time point, statistical significance was assessed using model-derived p values. To correct for multiple comparisons across time points and model terms, false discovery rate (FDR) correction was applied. An initial FDR correction was performed across all time points and the six effects tested in the omnibus model, using a significance level of q < 0.005. To reduce potentially spurious isolated transient detections, significant effects were required to extend across more than three consecutive time points (corresponding to more than 6 ms at the 500-Hz sampling rate). The time windows identified by this omnibus analysis were then used to constrain subsequent region-specific analyses. Within these time windows, FDR correction was performed separately for each recording region across the three effects tested (stimulus type, temporal condition, and their interaction), using q < 0.05.

To characterize the effects underlying significant region-specific model terms, we conducted post hoc comparisons at each time point. These comparisons identified whether responses differed between deviants and standards within each temporal condition, and whether responses to each stimulus type differed between rhythmic and jittered sequences. The post hoc models tested the overall deviant-versus-standard and rhythmic-versus-jittered effects, as well as four condition-specific pairwise comparisons: deviants versus standards during rhythmic sequences, deviants versus standards during jittered sequences, deviants during rhythmic versus jittered sequences, and standards during rhythmic versus jittered sequences. Each model included the corresponding contrast as a fixed effect and recording penetration as a random effect, with a random intercept and a random slope for the contrast to account for between-penetration variability in baseline responses and contrast effects. Post hoc p values were FDR corrected separately within each recording region across the tested contrasts and time points, using q < 0.05.

### Channel-wise response variability

To characterize response differences across recording channels and compartments, we calculated channel-wise response contrasts for stimulus type, temporal condition, and their interaction. To account for differences in baseline variability across channels, responses were normalized by the baseline standard deviation using the same procedure as described above. Three contrasts were calculated for each channel. The stimulus type contrast quantified the deviant–standard response difference averaged across temporal conditions, calculated as the mean of the deviant–standard differences under rhythmic and jittered stimulation. The temporal condition contrast quantified the difference between rhythmic and jittered responses, averaged across stimulus types. The stimulus type × temporal condition interaction contrast quantified the difference between the deviant–standard response differences in the rhythmic and jittered conditions. Positive interaction values indicated a larger deviant–standard difference under rhythmic stimulation, whereas negative values indicated a larger difference under jittered stimulation. These channel-wise contrasts were provided as descriptive measures of variability in response differences and were not interpreted as statistical significance measures.

### Directed functional connectivity analysis

Directed functional interactions between recording regions were assessed separately for LFP and AMUA using time-resolved Granger causality (Valt et al. 2024) as implemented in the Multivariate Granger Causality (MVGC) Toolbox for MATLAB (Barnett and Seth 2014). Analyses were performed within each penetration and separately for each combination of temporal condition (rhythmic vs. jittered) and stimulus type (deviant vs. standard). For each pair of recording regions, signals were analyzed in the 0–250 ms interval following stimulus onset. Granger causality was estimated using a multivariate autoregressive model with a maximum model order of 10 samples (20 ms), with the optimal model order determined using the Bayesian information criterion. Analyses were performed in sliding windows of 100 ms, advanced in 20 ms steps, yielding time-resolved estimates of directed interactions. Only region pairs with sufficient valid trials in both signals were included, and trials containing missing values in either recording region were excluded from the corresponding analysis.

For each directed connection, a directionality index (DI) was calculated from the pairwise-conditional Granger-causality estimates in the two directions. The DI was defined as the difference between the Granger-causality values in the two directions, divided by their sum: DI = (GC₁→₂ − GC₂→₁) / (GC₁→₂ + GC₂→₁ + ε), where ε is a small constant added to avoid division by zero. The index ranges approximately from −1 to +1, with positive values indicating stronger Granger-causal influence from region 1 to region 2 and negative values indicating stronger influence in the reverse direction. Values near zero indicate more balanced influence in the two directions. Thus, the DI captures the relative directionality of statistical interactions rather than their absolute strength.

Statistical significance of effects of temporal condition and stimulus deviance was assessed using 1,000 permutation-based surrogate estimates generated during the Granger-causality analysis. Effects of temporal condition and stimulus type on directionality were assessed by comparing DI values across conditions. The effect of temporal condition was quantified as the difference in DI between rhythmic and jittered sequences, collapsed across stimulus types; the effect of stimulus type was quantified as the difference between deviant and standard responses, collapsed across temporal conditions. The interaction was quantified as a difference-of-differences, testing whether the rhythmic–jittered difference in DI differed between deviants and standards. These contrasts tested whether the relative balance of directed statistical influence varied with temporal regularity, stimulus identity, or their interaction. For each directed connection and time window, two-sided permutation p values were calculated from the corresponding null distributions. These analyses were performed independently for each directed pair among the eight recording categories: primary superficial, middle (thalamorecipient), and deep cortex; non-primary superficial, middle (thalamorecipient), and deep cortex; and dorsal and ventral MGB.

## RESULTS

### Channel grouping

We first functionally assigned auditory cortical and MGB recording sites to one of eight recording categories (six cortical: superficial, middle or deep layers of A1 or non A1; and two thalamic: dorsal or ventral MGB), based on their response properties and stereotaxic positions. The six cortical sites included A1 (primary) and non-A1 (non-primary) regions of the auditory cortex, each divided into three laminar depths (superficial, middle (thalamorecipient), and deep). The two thalamic sites included ventral and dorsal MGB.

Cortical recordings were obtained from nine rats across 21 four-shank multielectrode penetrations, simultaneously with thalamic recordings (see Methods). Each rat had one chronic thalamic implant, which remained in place while several temporary cortical penetrations were made. Thus, thalamic recordings were obtained in each rat across multiple cortical penetration sites. The cortical penetrations were at right angles to the cortical surface, and therefore sampled 84 cortical columns. Cortical columns were classified as A1 or non-A1 based on their responses to white-noise and frequency-response-area stimuli, with A1 columns being characterized by consistent responses to pure tones and markedly shorter, more transient response durations (see Methods for details). Overall, 57% of columns were classified as A1 and 43% as non-A1. Recording sites within each column were further assigned to superficial, middle, or deep cortical layers based on the location and extent of the onset response CSD sink. Forty-two % of recording sites were classified as superficial, 21% as middle (thalamorecipient), and 37% as deep. This corresponded to 494 superficial electrodes (269 A1, 225 non-A1), 243 middle (thalamorecipient) electrodes (138 A1, 105 non-A1), and 437 deep sites (265 A1, 172 non-A1). MGB recording electrodes were subdivided into ventral and dorsal regions based on their position relative to the electrode with the strongest AMUA response (see Methods for details), yielding 111 ventral and 88 dorsal MGB channels.

### LFP responses

We first asked how temporal predictability shapes LFP responses across the auditory pathway, and whether its effects differ between anatomical regions or depend on stimulus identity. Having identified the eight recording categories spanning the auditory cortex and MGB, we examined how stimulus type (deviant vs. standard) and temporal condition (rhythm vs. jitter) modulated LFP responses across the sites. Time-resolved linear mixed-effects models revealed a significant main effect of temporal condition across several time windows following FDR correction (Figure 3A). Specifically, neural responses differed between rhythmic and jittered conditions from 26–42 ms, 58–96 ms, 120–184 ms, and 214–250 ms. In contrast, there was no significant main effect of stimulus type. Importantly, the effect of temporal condition varied significantly as a function of the recording site, with significant stimulus-by-site interactions observed during 4–44 ms, 48–98 ms, 102–186 ms, and 196–250 ms. In addition, a significant three-way interaction between stimulus type, temporal condition, and recording site was observed at 94–114 ms, 124–136 ms, and 168–176 ms. No significant interaction between stimulus type and temporal condition, or between stimulus type and recording site, was detected. All reported effects survived FDR correction and met the predefined minimum cluster-duration criterion.

**Figure 3.**
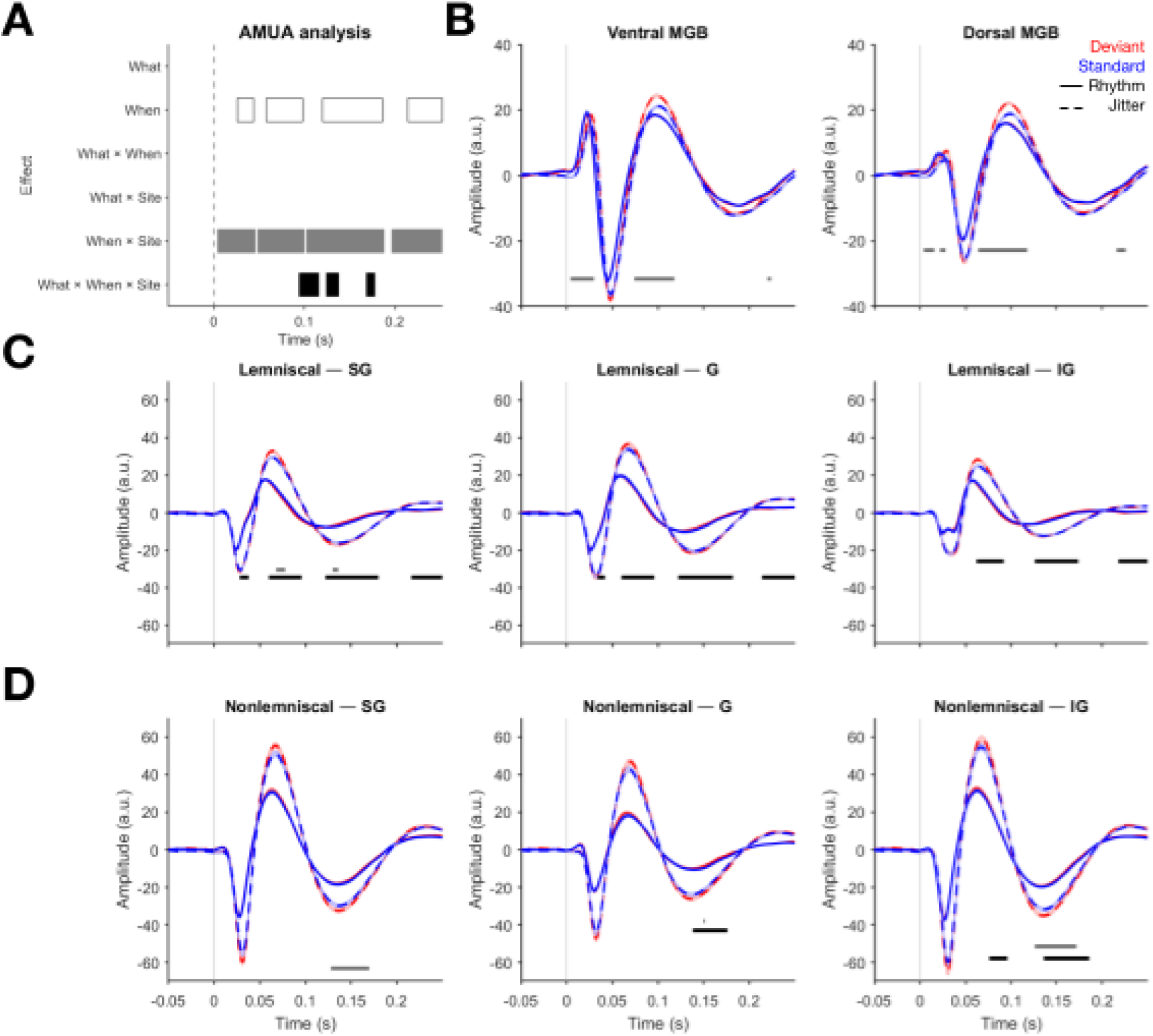
Effects of stimulus type and temporal condition on LFP signals across recording sites. **(A)** Post-stimulus-onset times at which significant main effects of stimulus type or temporal condition or recording site were observed. **(B)** Ventral and dorsal divisions of the MGB show significantly stronger responses in the jittered (temporally unpredictable) condition than in the rhythmic (temporally predictable) condition. Mean time-series are averaged over trials, channels and penetrations within each recording site. Shaded areas denote SEM over channels, pooled over all penetrations. Horizontal bars denote time windows of significant effects (p_FDR_ < 0.05). Amplitudes expressed as arbitrary units given normalization to pre-stimulus baseline (see Materials and methods), with values scaled by the standard deviation of the trial-averaged baseline activity for each channel. **(C)** Primary regions of the auditory cortex show significant interaction between stimulus type and temporal condition across cortical layers. Legend as in (B). **(D)** Non-primary regions of the auditory cortex show interactions between stimulus type and temporal condition across layers, as well as more confined differences between rhythmic and jittered conditions. Legend as in (B).

To determine where along the auditory pathway temporal predictability exerted its strongest effects, we examined the eight recording sites separately, including thalamic (Fig. 3B), primary auditory cortex (Fig. 3C), and non-primary auditory cortex (Fig. 3D). This allowed us to establish whether the effects observed across sites reflected a broadly shared cortical response or distinct patterns across anatomical subdivisions. To this end, we fitted separate time-resolved linear mixed-effects models within each site, to elucidate the two-way (temporal condition x recording site) and three-way (stimulus type x temporal condition x recording site) interactions identified above. The site-specific analyses revealed a broadly consistent pattern of temporal-predictability effects across the cortical recording sites. Significant effects were observed in superficial, middle (thalamorecipient), and deep layers of both primary and non-primary fields, with the most consistent effects occurring at approximately 60–96 ms and 120– 186 ms, together with shorter effects near stimulus onset and toward the end of the analyzed epoch (Table 1). The strongest effects were observed in the middle layers of the primary cortex and in deeper cortical layers, with peak statistics of *F*(1,10808) = 18.98 and *F*(1,10808) = 18.88, respectively. Post-hoc analysis revealed that, in all of the cortical recording sites, LFP amplitudes were stronger for stimuli presented in jittered than in rhythmic conditions (all p-values < 0.05, FDR-corrected).

**Table 1.** Site-specific effects of temporal condition on LFP amplitudes.

| **Recording site** | **Significant window (ms)** | **Peak F(1, df)** | **Peak p** | **Peak latency (ms)** |
|---|---|---|---|---|
| Lem-SG | 28–38 | 9.02 (10808) | .0027 | 32 |
|  | 60–96 | 16.41 (10808) | <.001 | 74 |
|  | 122–180 | 14.65 (10808) | <.001 | 146 |
|  | 216–250 | 10.77 (10808) | .0010 | 242 |
| Lem-G | 34–42 | 10.58 (10808) | .0012 | 38 |
|  | 60–96 | 18.98 (10808) | <.001 | 70 |
|  | 122–182 | 16.17 (10808) | <.001 | 138 |
|  | 214–250 | 11.33 (10808) | <.001 | 242 |
| Lem-IG | 62–92 | 18.88 (10808) | <.001 | 72 |
|  | 126–174 | 15.75 (10808) | <.001 | 144 |
|  | 218–250 | 10.28 (10808) | .0013 | 224 |
| NLem-SG | 32–38 | 7.33 (8264) | .0068 | 36 |
|  | 66–94 | 10.42 (8264) | .0013 | 84 |
|  | 122–186 | 12.53 (8264) | <.001 | 168 |
|  | 220–230 | 7.86 (8264) | .0051 | 224 |
| NLem-G | 138–176 | 11.09 (8264) | <.001 | 162 |
| NLem-IG | 76–96 | 9.84 (8264) | .0017 | 90 |
|  | 136–186 | 13.21 (8264) | <.001 | 176 |

Having established that temporal predictability robustly modulated cortical responses, we next asked whether it also influenced how the auditory system differentiated deviant from standard stimuli. These differences depended on both predictability and recording site, with significant effects observed in the MGB and predominantly in non-primary cortical sites (Table 2). In the cortical recording sites, these interactions occurred primarily at later latencies, with effects spanning 68–78 and 130–136 ms in primary superficial cortex and 128–172 ms in the non-primary superficial, middle, and deeper layers. Post-hoc comparisons indicated that these interactions were driven by a stronger deviant–standard difference under jittered stimulation. In the primary superficial cortex, the deviant–standard difference was significant under jittered stimulation at both 68–78 ms and 130–136 ms, but was not significant under rhythmic stimulation at either latency.

**Table 2.** Site-specific interaction effects of stimulus type and temporal condition on LFP amplitudes.

| **Recording site** | **Sig. window (ms)** | **Peak F(1, df)** | **Peak p** | **Peak latency (ms)** | **T, deviant > standard (rhythm)** | **p, deviant > standard (rhythm)** | **T, deviant > standard (jitter)** | **p, deviant > standard (jitter)** |
|---|---|---|---|---|---|---|---|---|
| Lem-SG | 68-78 | 8.61 (10808) | .0033 | 72 | 1.457 | .145 | 2.553 | .011* |
|  | 130-136 | 8.04 (10808) | .0045 | 134 | .729 | .466 | 3.547 | <.001* |
| NLem-SG | 128-170 | 10.04 (8264) | .0015 | 152 | -.224 | .823 | 2.519 | .012* |
| NLem-G | 150-152 | 9.03 (8264) | .0026 | 150 | .416 | .678 | 2.578 | .010* |
| NLem-IG | 126-172 | 10.86 (8264) | <.001 | 148 | -.141 | .888 | 2.477 | .013* |
| MGB-D | 4-16 | 9.88 (13352) | .0016 | 4 | -.404 | .686 | -1.371 | .170 |
|  | 22-28 | 10.737 (13352) | .0011 | 6 | -.089 | .929 | 1.357 | .175 |
|  | 64-118 | 17.433 (13352) | <.001 | 54 | .331 | .741 | 3.947 | <.001* |
|  | 216-226 | 10.376 (13352) | .0012 | 222 | 1.721 | .085 | -2.456 | .014* |
| MGB-V | 4-30 | 17.335 (13352) | <.001 | 26 | -.903 | .366 | 1.565 | .118 |
|  | 74-118 | 15.214 (13352) | <.001 | 104 | -.630 | .529 | 3.649 | <.001* |
|  | 220-224 | 8.518 (13352) | .0035 | 222 | 2.367 | .018* | -1.194 | .233 |

In the MGB, stimulus type × temporal condition interactions were evident at early and late latencies, spanning 4–118 ms and again around 216–226 ms. During the early response, significant deviant–standard differences under jittered stimulation were observed from 64–118 ms in dorsal MGB and from 74–118 ms in ventral MGB. The shorter-latency comparisons (22– 28 ms in dorsal MGB and 4–30 ms in ventral MGB) were no longer significant in the revised post hoc models. At later latencies, the pattern of significant simple effects differed between the two subdivisions. In dorsal MGB, deviant responses were significantly weaker than standard responses during jittered stimulation, whereas the deviant–standard difference under rhythmic stimulation was not significant. In ventral MGB, deviant responses were significantly stronger than standard responses during rhythmic stimulation, whereas the difference under jittered stimulation was not significant.

To visualize how stimulus type and temporal conditions jointly shaped auditory responses at a single-channel level across recording compartments, we calculated channel-level contrasts for stimulus type (deviant vs. standard), temporal condition (rhythmic vs. jittered), and their interaction. Based on the results reported in Fig. 3, contrasts were calculated separately for LFP responses at 60–100 ms (Fig. 5A) and 120–190 ms (Fig. 5). These visualizations provide a complementary summary of the variability of response patterns across compartments.

Together, these findings indicate that temporal predictability shapes LFP responses across the auditory pathway, with jittered stimulation generally eliciting stronger responses than rhythmic stimulation. The influence of temporal predictability on deviant–standard differentiation depended on anatomical location and latency. In cortex and during intermediate-latency MGB responses, significant deviant–standard differences were predominantly observed under jittered stimulation. At later MGB latencies, the significant deviant–standard difference was observed under rhythmic stimulation in ventral MGB and under jittered stimulation in dorsal MGB, although the corresponding simple effect in the other temporal condition was not significant in either subdivision.

### AMUA responses

We next asked whether the effects of temporal predictability observed in LFP responses were also evident in AMUA, and whether their anatomical distribution was similar across the two measures. Across recording sites, time-resolved linear mixed-effects models revealed a significant main effect of temporal condition from 30–70 ms following FDR correction (Fig. 4A). The effect of temporal condition also varied across recording sites, with a significant temporal condition × recording site interaction extending from 10–214 ms. All reported effects survived FDR correction and the predefined minimum cluster-duration criterion.

**Figure 4.**
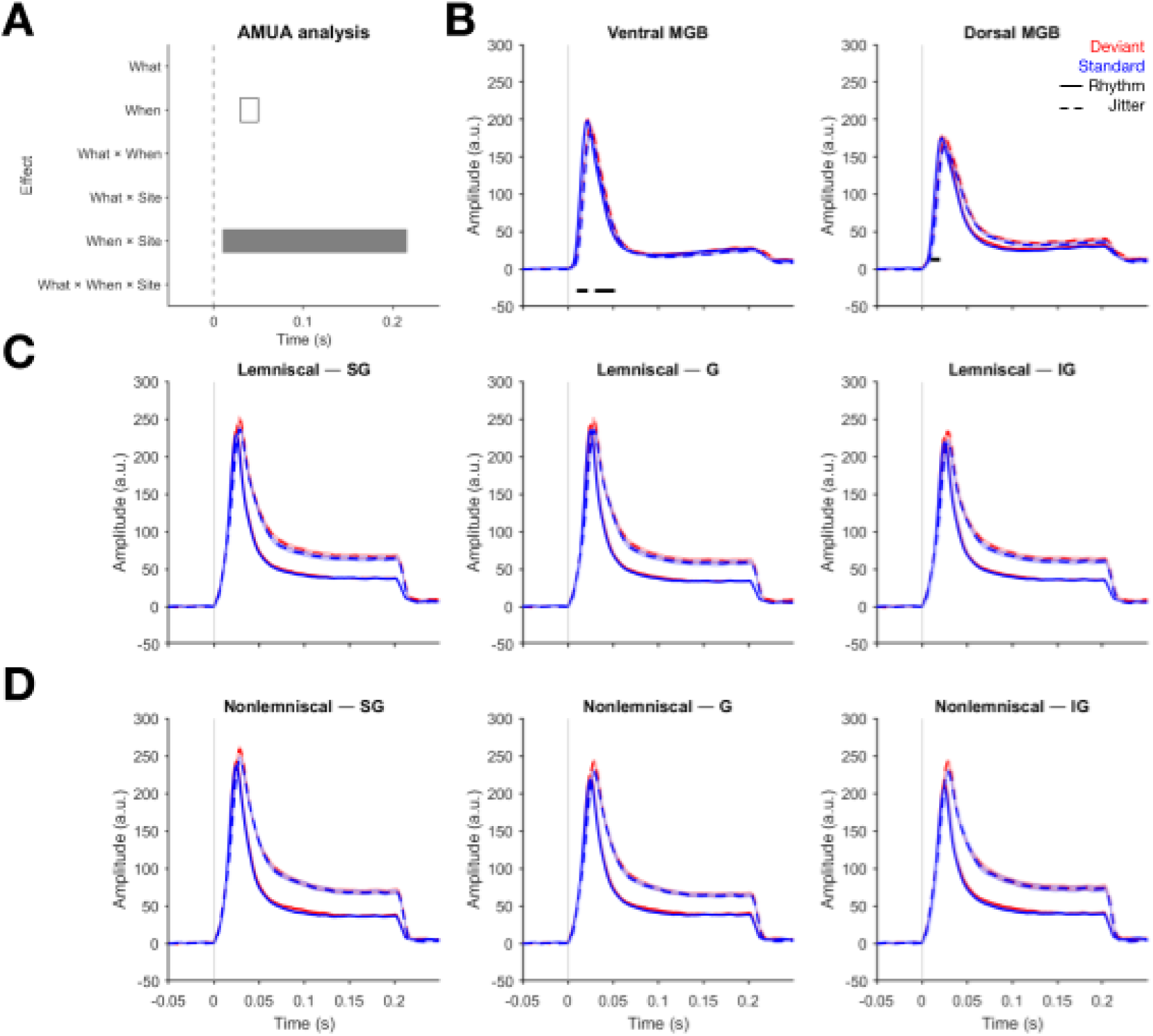
Effects of stimulus type and temporal condition on AMUA signals across recording sites. **(A)** Significant effects of stimulus type, temporal condition, and recording site. **(B)** Ventral and dorsal divisions of the MGB show significantly stronger responses in the jittered (temporally unpredictable) condition than in the rhythmic (temporally predictable) condition. Mean time-series are averaged over trials, channels and penetrations within each recording site. Shaded areas denote SEM over channels, pooled over all penetrations. Horizontal bars denote time windows of significant effects (p_FDR_ < 0.05). Amplitudes expressed as arbitrary units given normalization to pre-stimulus baseline (see Materials and methods), with values scaled by the standard deviation of the trial-averaged baseline activity for each channel. **(C)** Primary regions of the auditory cortex. Legend as in (B). **(D)** Non-primary regions of the auditory cortex. Legend as in (B).

To determine where these effects arose, we examined the individual anatomical compartments. Significant temporal-predictability effects were confined to the MGB (Fig. 4B), with no significant effects detected at the cortical recording sites after FDR correction (Fig. 4C–D). In dorsal MGB, a significant effect was observed from 10–20 ms, peaking at 16 ms (*F*(1,13352) = 33.38, *p* < .001), while in ventral MGB, significant effects occurred from 10–22 ms and 30– 52 ms, with peak statistics of *F*(1,13352) = 71.38 and *F*(1,13352) = 14.99, respectively (all *p*s < .001). Post-hoc analysis revealed that, in all significant time windows, AMUA amplitudes were stronger for stimuli presented in jittered than in rhythmic conditions (all p-values < 0.05, FDR-corrected). Response variability across channels, averaged within the 10-50 ms time window, is visualized in Fig. 5C.

**Figure 5.**
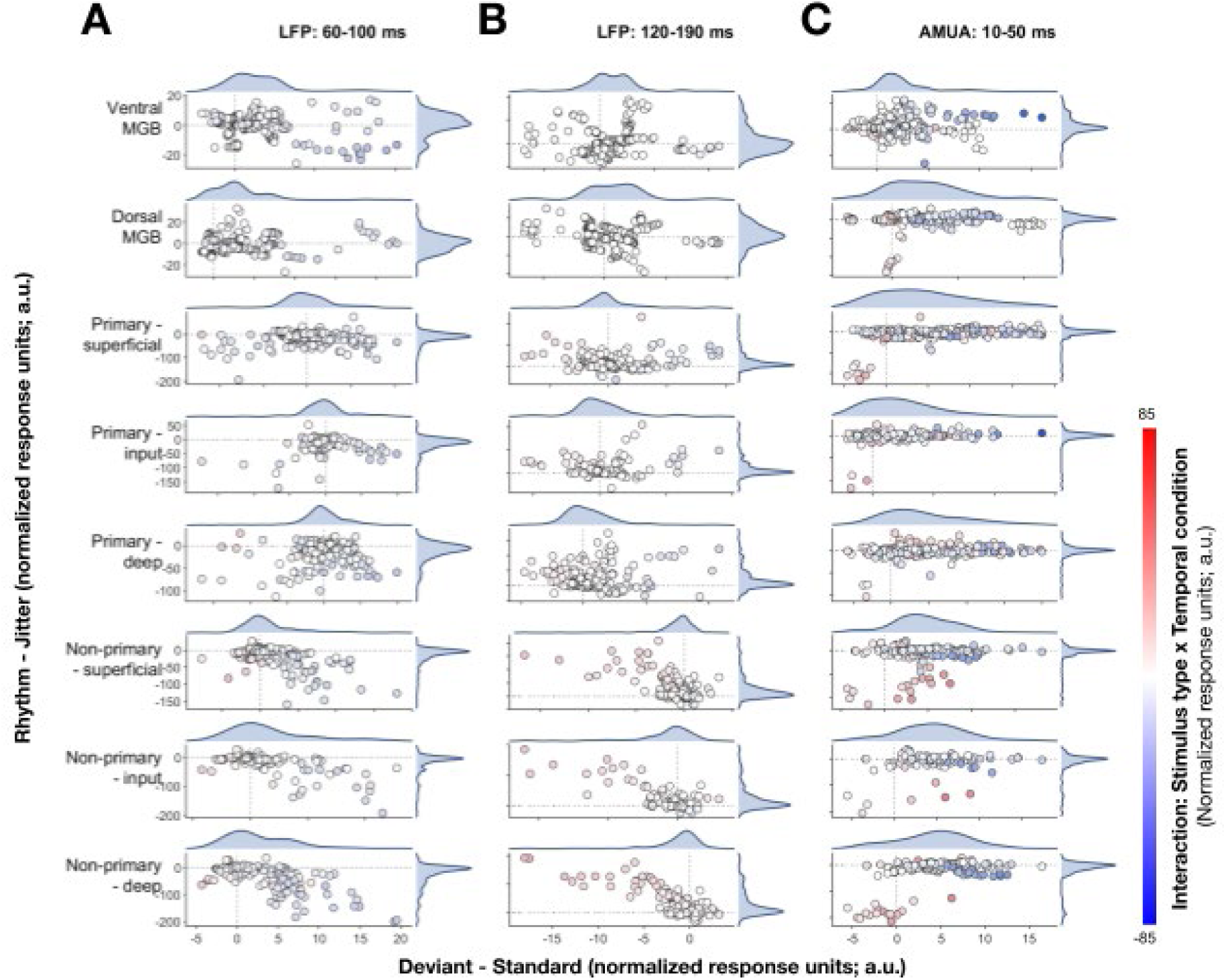
Channel-level auditory response contrasts. Scatterplots show stimulus type effects (X-axis) and temporal condition effects (Y-axis) for eight groups of electrodes across three analyses: (A) LFP responses at 60–100 ms, (B) LFP responses at 120–190 ms, and (C) AMUA responses at 10–50 ms. Each point is a single channel. Rows show distinct groups of electrodes. Responses were normalized by the pooled baseline standard deviation (see Methods). The stimulus type contrast represents the average deviant-versus-standard response difference across rhythmic and jittered conditions. The temporal condition contrast represents the average response in the rhythmic condition minus that in the jittered condition; positive values indicate larger responses in the rhythmic condition, whereas negative values indicate larger responses in the jittered condition. Point color represents the stimulus type × temporal condition interaction, reflecting the difference in deviant-versus-standard response contrasts between temporal conditions (red: deviant–standard response difference is larger in the rhythmic condition than in the jittered condition; blue deviant–standard response difference is larger in the jittered condition than in the rhythmic condition). Marginal density plots show the distributions of stimulus identity and predictability contrasts within each compartment and analysis.

### Directed functional connectivity analysis

The local response analyses revealed that temporal predictability modulated neural activity across the auditory pathway, but they could not establish whether it also altered the directed interactions between thalamic and cortical populations. We therefore used pairwise Granger causality in sliding time windows to test whether directed functional connectivity differed between rhythmic and jittered stimulation, and whether these differences depended on stimulus identity. In the analysis of LFP signals, no directed connectivity effects - i.e., differences in directionality index (DI) values between experimental conditions - survived FDR correction. At the uncorrected level, several jitter-specific directed interactions were observed, predominantly during the 52–172 ms post-stimulus period. These included stronger directed influences in the jittered condition from non-primary superficial and deep sites toward primary middle (thalamorecipient) and deep sites, as well as interactions within the primary sites. Given that none of these effects survived correction for multiple comparisons, they are not considered statistically significant; the uncorrected results are shown for completeness in Fig. 6A.

**Figure 6.**
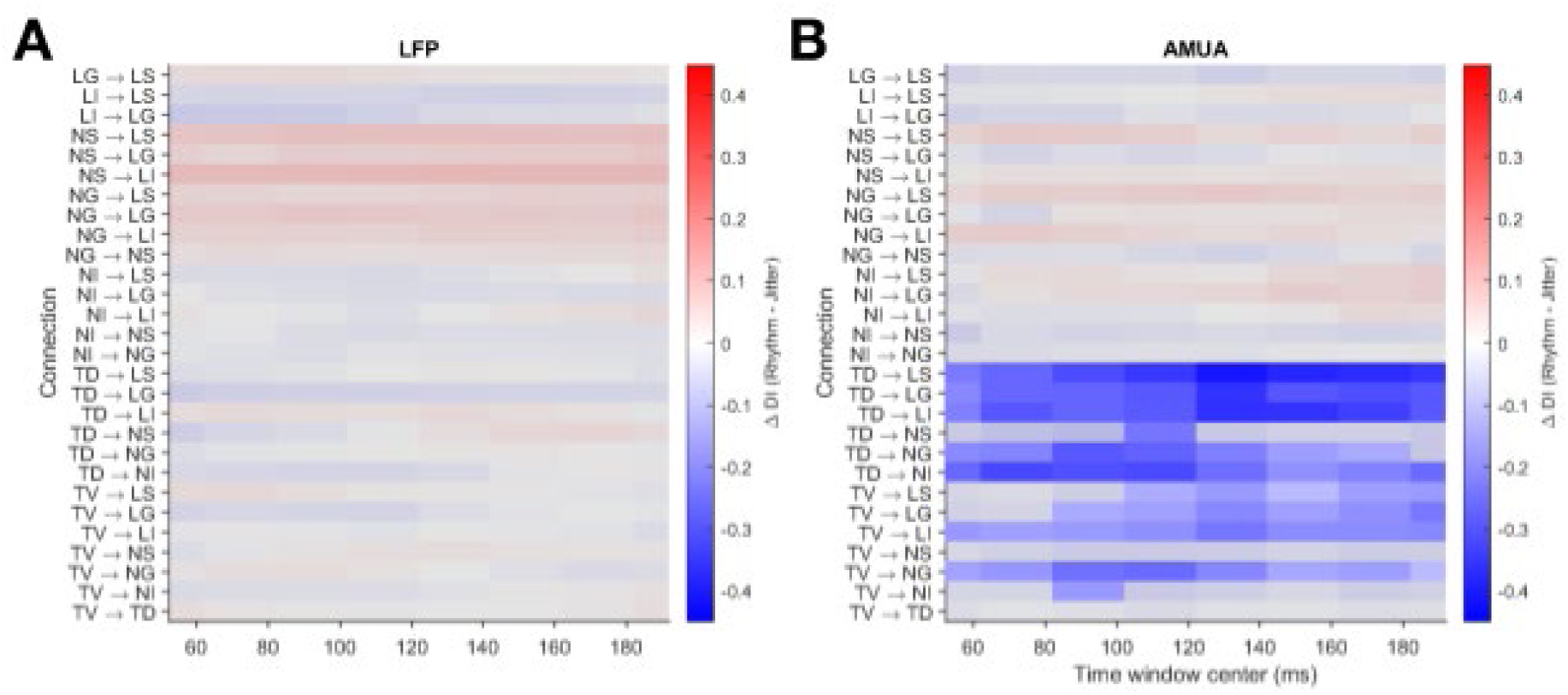
Effects of temporal condition on pairwise directionality index (DI) values. **(A)** Results based on LFP signals. No significant effects were found. High vs. low values (red vs. blue) denote stronger vs. weaker directed functional connectivity between regions in the rhythmic compared to jittered condition. P: primary, N: non-primary, S: superficial, G: middle (thalamorecipient), I: deep, T: thalamus (MGB), D: dorsal, V: ventral. (B) Results based on AMUA signals. Connectivity from (primarily dorsal) MGB to both primary and non-primary auditory cortical regions is more pronounced in the jittered compared to rhythmic temporal condition conditions. Unmasked area denotes p_FDR_<0.05.

Whereas the LFP analysis revealed no significant condition-dependent differences in directed connectivity after FDR correction, the AMUA analysis (Fig. 6B) revealed robust differences between rhythmic and jittered stimulation. Significant directed influences originated from both dorsal and ventral MGB, and were directed toward cortical recording sites, including primary and non-primary superficial, middle (thalamorecipient), and deep layers. These effects were significant between approximately 52 and 172 ms, with particularly widespread connectivity from dorsal MGB to cortical sites and a similar pattern from ventral MGB. Together, these results suggest that temporal predictability is associated not only with changes in local response magnitude but also with changes in directed functional interactions between thalamic and cortical populations. These effects were evident in AMUA, where jittered stimulation was associated with stronger directed influences from the MGB toward distributed cortical sites, but were not significant in the LFP analysis after correction for multiple comparisons. Granger causality provides evidence of statistical directionality, however, and does not by itself establish direct causal information transfer.

## DISCUSSION

This study asked whether regularities in sound identity (’what’) and stimulus timing (’when’) are represented in dissociable parts of the auditory thalamocortical system. Three results stand out. First, rhythmic temporal structure suppressed LFP responses broadly, across primary and non-primary cortex and across cortical layers. Second, sound identity produced only weak, context-dependent deviant–standard effects, emerging mainly as an interaction with temporal condition rather than as a stable main effect. Third, temporal condition modulated AMUA and directed functional interactions primarily at the thalamic level, with jitter-specific MGB-to-cortex influences extending across cortical fields and layers. Together, these findings identify temporal regularity as a pervasive modulator of auditory thalamocortical processing, while showing that identity-related responses depend on temporal context, anatomical location, and response latency. Although temporal regularity modulated cortical LFP responses across all sampled fields and layers, its effects on AMUA were confined to the MGB, and deviance-related effects varied across cortical fields and MGB subdivisions. In addition, jitter-specific directed statistical influences from MGB to cortex were evident in AMUA but not in LFP after correction for multiple comparisons. These findings argue against a simple one-to-one mapping between temporal-context processing and a single anatomical subdivision, while indicating that the expression of temporal and identity-related effects varies across levels of the auditory thalamocortical system (Rauschecker and Tian 2000; Auksztulewicz et al. 2018; Cappotto et al. 2023; Auksztulewicz et al. 2025; Ten Oever and Martin 2024).

### Evoked responses are smaller for rhythmic rather than jittered sequences

The most robust and consistent effect was larger cortical LFP and thalamic AMUA responses to jittered than to rhythmic stimulation. This may seem counterintuitive if ‘when’ predictability is assumed to enhance processing through anticipatory alignment of neural excitability (Lakatos et al. 2007, 2013; Schroeder and Lakatos 2009). However, the temporal-expectation literature does not predict uniform enhancement: auditory N1-like responses have been reported to increase, decrease, or remain unchanged under ‘when’ predictability, depending on whether expectation is coupled to attention, task relevance, or motor preparation (Nobre and van Ede 2017; Breska and Deouell 2017). Our data instead align with a regularity-related suppression account. In human EEG and subcortical recordings, temporally predictable sequences enhance repetition suppression, while jitter reduces the sensory-memory signatures associated with regularity encoding (Garrido, Kilner, Kiebel, et al. 2009; Heilbron and Chait 2018). The present results extend previous observations of regularity-related suppression by characterizing its distribution across cortical layers, auditory cortical fields, and thalamic subdivisions. Rhythmic stimulation was associated with smaller LFP responses across all sampled cortical layers in both A1 and non-A1 cortex, with the most consistent effects occurring at intermediate poststimulus latencies and the strongest statistical effects in middle-layer A1 and deep cortical sites. In contrast, jitter-related increases in AMUA were detected in both dorsal and ventral MGB, beginning at approximately 10 ms after stimulus onset, but were not significant in cortical recordings after correction for multiple comparisons. Thus, temporal-context modulation was widespread but was expressed differently across recording sites and physiological measures. These differences suggest that the effects of temporal regularity cannot be described simply as a uniform change in response magnitude throughout the auditory pathway; rather, their expression depends on the neural population and signal being measured.

Mechanistically, a rhythmic sequence may allow the auditory system to build a temporally structured model of the sequence - analogous to the hierarchical sequence representations proposed for transition-probability and pattern learning more broadly (Dehaene et al. 2015) - reducing the response needed when an expected sound arrives at an expected time; the larger jitter response would then reflect weaker temporal prediction or regularity encoding. This need not imply worse processing of rhythmic sounds, as a smaller population response can coexist with more efficient encoding or more precise representations (Kok et al. 2012; Chalk et al. 2018). Repetition-related suppression may also contribute to the observed effect. Regular inter-stimulus spacing could strengthen response suppression through short-term adaptation, which is sensitive to stimulus history and temporal structure (Ulanovsky et al. 2004; Carbajal and Malmierca 2018). However, the present results do not establish the contribution of adaptation relative to temporal prediction or other forms of context-dependent response modulation. These mechanisms may also operate together. We therefore interpret the larger responses to jittered sequences as evidence of temporal-context-dependent modulation, without assigning the effect to a specific underlying mechanism. What the data do establish is that the effect was not confined to a specialized cortical network: suppression was present in middle (thalamorecipient), superficial, and deep layers of both A1 and non-A1 cortex, and appeared at short latency in both divisions of MGB (from ∼10 ms post-onset), arguing for a system-wide, rather than localized, influence of temporal context.

### Identity effects depend on temporal context

Identity effects became apparent once temporal condition was considered explicitly. In cortex, deviant–standard differences were generally larger under jitter, especially at later latencies and most consistently in non-primary fields; the same pattern emerged in early MGB windows. This is consistent with a growing literature showing that ‘what’ and ‘when’ predictions interact rather than combine additively - for example, suppression to an auditory event is strongest when both its timing and identity are predictable, and is reduced when either dimension is uncertain (Auksztulewicz et al. 2018; Ten Oever and Martin 2024).

Two (non-exclusive) accounts can explain why deviance sensitivity was *larger* under jitter. First, rhythmic timing may induce a general suppressive state that attenuates both standards and deviants once onset time is highly expected, while jitter - by weakening this global suppression - preserves sensitivity to identity differences. Second, when onset time is uncertain, the occurrence of a sound may itself carry more information, causing identity to be weighted more heavily in updating the inferred sequence state; this second account fits the predominance of later, integrative interaction effects in non-primary cortex, which is well positioned to incorporate context over longer windows.

A further feature of the data was a late effect (∼216–226 ms) in both MGB subdivisions, although the pattern of significant simple effects differed between them. In dorsal MGB, deviants elicited weaker responses than standards under jittered stimulation, whereas the deviant–standard difference under rhythmic stimulation did not reach significance. In ventral MGB, deviants elicited stronger responses than standards under rhythmic stimulation, whereas the difference under jittered stimulation was not significant. These findings suggest that late deviant–standard differentiation may depend on both temporal context and MGB subdivision, although the revised post hoc analyses do not establish a significant crossover within either subdivision. One possible explanation is temporally evolving, feedback-dependent reweighting of identity information (Bartlett 2013; Kommajosyula et al. 2021) - plausibly via corticothalamic feedback, given the latency and the fact that both MGB divisions are targets of extensive descending projections (Pandya et al. 1994; Kimura et al. 2003; E. G. Jones 2001; Lakatos et al. 2020). We present this as a candidate signature of temporally evolving context-dependence rather than evidence for a specific computational mechanism.

### Weak deviance effects reflect context-dependence, not absence of regularity encoding

Unlike classic oddball studies, we did not observe a robust, context-independent deviant > standard effect. This is an important qualification of the conventional mismatch framework rather than evidence against predictive or regularity-based processing. A standard–deviant contrast conflates several factors - repetition history, frequency-specific adaptation, transition probability, and stimulus rarity - that can reproduce mismatch-like effects through local synaptic depression alone, without requiring a distinct prediction-error signal (Näätänen et al. 2007; Garrido, Kilner, Stephan, et al. 2009; Antunes et al. 2010). Because ‘what’ prediction (stimulus type: deviant vs. standard) and ‘when’ predictability (temporal condition: rhythmic vs. jittered) were manipulated orthogonally here, our design reveals that deviance responses are themselves conditional on temporal context, rather than reflecting a single generic computation - consistent with evidence that even pattern-sensitive, context-dependent deviance coding in the auditory midbrain is not reducible to simple adaptation (Malmierca et al. 2019). Pooling across heterogeneous, differently-tuned recording sites, and the use of an anesthetized preparation - which can reduce arousal-related gain and weaken long-range recurrent and corticothalamic feedback - likely also diluted any local deviance signal, particularly for later, contextually-integrated components (Lakatos et al. 2020).

### MGB as a hub for temporal-context modulation

The AMUA and directed-connectivity (Granger causality) results converge on MGB as a prominent site of temporal-context modulation. Temporal jitter increased AMUA in both dorsal and ventral MGB from short latency, before the later cortical identity × timing interactions emerged, and jitter-specific directed influences ran predominantly from MGB to cortex (∼52–172 ms), spanning primary and non-primary fields and superficial, middle (thalamorecipient), and deep layers alike. This argues against a strict division in which ventral (lemniscal) MGB carries only acoustic content while dorsal (nonlemniscal) MGB carries only temporal or contextual information - a division that is well established anatomically and physiologically (Calford 1983; Anderson et al. 2007; de Venecia et al. 1995; Jones 1998; Jones 2003) but does not appear to hold functionally here. Instead, ventral MGB appears to contribute temporally structured sensory drive to core cortex, consistent with its role in relaying fine temporal structure such as musical rhythm and tone (Musacchia et al. 2014), while dorsal MGB contributes more broadly distributed contextual/gain-related influences, in keeping with its diffuse, patchy corticothalamic projection pattern (McMullen and de Venecia 1993; Hashikawa et al. 1995). We are cautious in interpreting these directed measures: Granger causality indexes directed statistical dependence, not monosynaptic or anatomically verified causal transmission, and unrecorded common inputs or corticothalamic loops could contribute to the apparent directionality. Notably, this jitter-specific directionality was robust in AMUA but did not survive correction in LFP, suggesting the effect is more reliably expressed in population spiking than in the aggregate synaptic/field signal.

### Implications, caveats, and future directions

These findings support a model in which auditory identity and temporal regularity are integrated across the thalamocortical system rather than allocated to segregated ‘what’ and ‘when’ pathways, with neither primary/non-primary anatomy nor MGB subdivision mapping cleanly onto this functional distinction. The pattern is compatible with predictive-processing accounts but does not uniquely support them over adaptation-, gain-, or state-based mechanisms, and we do not claim to adjudicate between these here. Because recordings were made under anesthesia, our results speak to relatively automatic components of temporal-regularity processing; the presence of robust effects under anesthesia is itself informative, but the strength and even direction of these effects may not generalize directly to awake, attentive states, where arousal, task demands, and corticothalamic feedback are known to shape temporal-expectation effects (Lakatos et al. 2013; Nobre and van Ede 2017). Future work should (1) dissociate repetition, transition probability, and ‘when’ predictability more cleanly - e.g., with many-standards, cascade, and omission controls under independently manipulated inter-stimulus-interval distributions; (2) record in awake, behaving animals; and (3) combine simultaneous IC–MGB–cortex recording with perturbation of corticothalamic feedback to test whether early MGB effects are inherited, locally generated, or top-down in origin, and whether dorsal and ventral MGB make dissociable causal contributions to temporal gain control and deviance processing.

## Conclusion

Rhythmic temporal regularity suppressed cortical LFP responses and reduced early MGB AMUA relative to jittered stimulation, consistent with regularity-related predictive or adaptive suppression. Identity deviance did not produce a uniform mismatch response; instead, deviant– standard effects were conditional on temporal context, emerging primarily in MGB and later cortical responses, with a late reversal in thalamus. Temporal context is thus a pervasive organizing dimension of auditory thalamocortical processing, shaping local response amplitude and directed thalamocortical interactions, whereas stimulus-identity effects depend on temporal context.

## ACKNOWLEDGMENTS

We would like to thank HiJee Kang for assistance with setting up the experiment.

## REFERENCES

An, Hyunjung, Ryszard Auksztulewicz, Hijee Kang, and Jan W. H. Schnupp. 2021. “Cortical Mapping of Mismatch Responses to Independent Acoustic Features.” Hearing Research 399 (107894): 107894.

Anderson, L. A., M. N. Wallace, and A. R. Palmer. 2007. “Identification of Subdivisions in the Medial Geniculate Body of the Guinea Pig.” Hearing Research 228 (1–2): 156–67.

Anderson, Lucy A., and Jennifer F. Linden. 2011. “Physiological Differences between Histologically Defined Subdivisions in the Mouse Auditory Thalamus.” Hearing Research 274 (1–2): 48–60.

Antunes, Flora M., Israel Nelken, Ellen Covey, and Manuel S. Malmierca. 2010. “Stimulus-Specific Adaptation in the Auditory Thalamus of the Anesthetized Rat.” PloS One 5 (11): e14071.

Auksztulewicz, Ryszard, Ozan Bahattin Ödül, Saskia Helbling, et al. 2025. “‘What’ and ‘When’ Predictions Jointly Modulate Speech Processing.” The Journal of Neuroscience: The Official Journal of the Society for Neuroscience 45 (20). 10.1523/JNEUROSCI.1049-24.2025.

Auksztulewicz, Ryszard, Caspar M. Schwiedrzik, Thomas Thesen, et al. 2018. “Not All Predictions Are Equal: ‘What’ and ‘When’ Predictions Modulate Activity in Auditory Cortex through Different Mechanisms.” The Journal of Neuroscience: The Official Journal of the Society for Neuroscience 38 (40): 8680–93.

Barnett, Lionel, and Anil K. Seth. 2014. “The MVGC Multivariate Granger Causality Toolbox: A New Approach to Granger-Causal Inference.” Journal of Neuroscience Methods 223 (February): 50–68.

Bartlett, Edward L. 2013. “The Organization and Physiology of the Auditory Thalamus and Its Role in Processing Acoustic Features Important for Speech Perception.” Brain and Language 126 (1): 29–48.

Bizley, Jennifer K., Fernando R. Nodal, Israel Nelken, and Andrew J. King. 2005. “Functional Organization of Ferret Auditory Cortex.” Cerebral Cortex (New York, N.Y.: 1991) 15 (10): 1637–53.

Breska, Assaf, and Leon Y. Deouell. 2017. “Neural Mechanisms of Rhythm-Based Temporal Prediction: Delta Phase-Locking Reflects Temporal Predictability but Not Rhythmic Entrainment.” PLOS Biology 15 (2): e2001665.

Calford, M. B. 1983. “The Parcellation of the Medial Geniculate Body of the Cat Defined by the Auditory Response Properties of Single Units.” The Journal of Neuroscience: The Official Journal of the Society for Neuroscience 3 (11): 2350–64.

Cappotto, Drew, Dan Luo, Hiu Wai Lai, et al. 2023. “‘What’ and ‘When’ Predictions Modulate Auditory Processing in a Mutually Congruent Manner.” Frontiers in Neuroscience 17 (September): 1180066.

Carbajal, Guillermo V., and Manuel S. Malmierca. 2018. “The Neuronal Basis of Predictive Coding Along the Auditory Pathway: From the Subcortical Roots to Cortical Deviance Detection.” Trends in Hearing 22: 2331216518784822.

Chalk, Matthew, Olivier Marre, and Gašper Tkačik. 2018. “Toward a Unified Theory of Efficient, Predictive, and Sparse Coding.” Proceedings of the National Academy of Sciences of the United States of America 115 (1): 186–91.

Cheveigné, Alain de. 2020. “ZapLine: A Simple and Effective Method to Remove Power Line Artifacts.” NeuroImage 207 (116356): 116356.

De Cheveigné, Alain, and Jonathan Z. Simon. 2008. “Denoising Based on Spatial Filtering.” Journal of Neuroscience Methods 171 (2): 331–39.

Dehaene, Stanislas, Florent Meyniel, Catherine Wacongne, Liping Wang, and Christophe Pallier. 2015. “The Neural Representation of Sequences: From Transition Probabilities to Algebraic Patterns and Linguistic Trees.” Neuron 88 (1): 2–19.

Ferro, Demetrio, Jochem van Kempen, Michael Boyd, Stefano Panzeri, and Alexander Thiele. 2021. “Directed Information Exchange between Cortical Layers in Macaque V1 and V4 and Its Modulation by Selective Attention.” Proceedings of the National Academy of Sciences of the United States of America 118 (12): e2022097118.

Friston, Karl. 2005. “A Theory of Cortical Responses.” Philosophical Transactions of the Royal Society of London. Series B, Biological Sciences 360 (1456): 815–36.

Garrido, Marta I., James M. Kilner, Stefan J. Kiebel, Klaas E. Stephan, Torsten Baldeweg, and Karl J. Friston. 2009. “Repetition Suppression and Plasticity in the Human Brain.” NeuroImage 48 (1): 269–79.

Garrido, Marta I., James M. Kilner, Klaas E. Stephan, and Karl J. Friston. 2009. “The Mismatch Negativity: A Review of Underlying Mechanisms.” Clinical Neurophysiology: Official Journal of the International Federation of Clinical Neurophysiology 120 (3): 453–63.

Happel, Max F. K., Marcus Jeschke, and Frank W. Ohl. 2010. “Spectral Integration in Primary Auditory Cortex Attributable to Temporally Precise Convergence of Thalamocortical and Intracortical Input.” The Journal of Neuroscience: The Official Journal of the Society for Neuroscience 30 (33): 11114–27.

Hashikawa, T., M. Molinari, E. Rausell, and E. G. Jones. 1995. “Patchy and Laminar Terminations of Medial Geniculate Axons in Monkey Auditory Cortex.” The Journal of Comparative Neurology 362 (2): 195– 208.

Heilbron, Micha, and Maria Chait. 2018. “Great Expectations: Is There Evidence for Predictive Coding in Auditory Cortex?” Neuroscience 389 (October): 54–73.

Ives, D. Timothy, David R. R. Smith, and Roy D. Patterson. 2005. “Discrimination of Speaker Size from Syllable Phrases.” The Journal of the Acoustical Society of America 118 (6): 3816–22.

Jaramillo, Santiago, and Anthony M. Zador. 2011. “The Auditory Cortex Mediates the Perceptual Effects of Acoustic Temporal Expectation.” Nature Neuroscience 14 (2): 246–51.

Jones, E. G. 1998. “A New View of Specific and Nonspecific Thalamocortical Connections.” Advances in Neurology 77: 49–71; discussion 72-3.

Jones, E. G. 2001. “The Thalamic Matrix and Thalamocortical Synchrony.” Trends in Neurosciences 24 (10): 595–601.

Jones, Edward G. 2003. “Chemically Defined Parallel Pathways in the Monkey Auditory System.” Annals of the New York Academy of Sciences 999 (1): 218–33.

Kajikawa, Yoshinao, Lisa de La Mothe, Suzanne Blumell, and Troy A. Hackett. 2005. “A Comparison of Neuron Response Properties in Areas A1 and CM of the Marmoset Monkey Auditory Cortex: Tones and Broadband Noise.” Journal of Neurophysiology 93 (1): 22–34.

Kawahara, Hideki. 2006. “STRAIGHT, Exploitation of the Other Aspect of VOCODER: Perceptually Isomorphic Decomposition of Speech Sounds.” Acoustical Science and Technology 27 (6): 349–53.

Kimura, A., T. Donishi, T. Sakoda, M. Hazama, and Y. Tamai. 2003. “Auditory Thalamic Nuclei Projections to the Temporal Cortex in the Rat.” Neuroscience 117 (4): 1003–16.

Kok, Peter, Janneke F. M. Jehee, and Floris P. de Lange. 2012. “Less Is More: Expectation Sharpens Representations in the Primary Visual Cortex.” Neuron 75 (2): 265–70.

Kommajosyula, Srinivasa P., Edward L. Bartlett, Rui Cai, Lynne Ling, and Donald M. Caspary. 2021. “Corticothalamic Projections Deliver Enhanced Responses to Medial Geniculate Body as a Function of the Temporal Reliability of the Stimulus.” The Journal of Physiology 599 (24): 5465–84.

Lakatos, Peter, Chi-Ming Chen, Monica N. O’Connell, Aimee Mills, and Charles E. Schroeder. 2007. “Neuronal Oscillations and Multisensory Interaction in Primary Auditory Cortex.” Neuron 53 (2): 279–92.

Lakatos, Peter, Gabriella Musacchia, Monica N. O’Connel, Arnaud Y. Falchier, Daniel C. Javitt, and Charles E. Schroeder. 2013. “The Spectrotemporal Filter Mechanism of Auditory Selective Attention.” Neuron 77 (4): 750–61.

Lakatos, Peter, Monica N. O’Connell, Annamaria Barczak, et al. 2020. “The Thalamocortical Circuit of Auditory Mismatch Negativity.” Biological Psychiatry 87 (8): 770–80.

Lee, Charles C., and Jeffery A. Winer. 2008. “Connections of Cat Auditory Cortex: I. Thalamocortical System.” The Journal of Comparative Neurology 507 (6): 1879–900.

Lomber, Stephen G., and Shveta Malhotra. 2008. “Double Dissociation of ‘what’ and ‘Where’ Processing in Auditory Cortex.” Nature Neuroscience 11 (5): 609–16.

Malmierca, Manuel S., Blanca E. Niño-Aguillón, Javier Nieto-Diego, Ángel Porteros, David Pérez-González, and Carles Escera. 2019. “Pattern-Sensitive Neurons Reveal Encoding of Complex Auditory Regularities in the Rat Inferior Colliculus.” NeuroImage 184 (January): 889–900.

McMullen, N. T., and R. K. de Venecia. 1993. “Thalamocortical Patches in Auditory Neocortex.” Brain Research 620 (2): 317–22.

Musacchia, Gabriella, Edward W. Large, and Charles E. Schroeder. 2014. “Thalamocortical Mechanisms for Integrating Musical Tone and Rhythm.” Hearing Research 308 (February): 50–59.

Näätänen, R., P. Paavilainen, T. Rinne, and K. Alho. 2007. “The Mismatch Negativity (MMN) in Basic Research of Central Auditory Processing: A Review.” Clinical Neurophysiology: Official Journal of the International Federation of Clinical Neurophysiology 118 (12): 2544–90.

Nobre, Anna C., and Freek van Ede. 2017. “Anticipated Moments: Temporal Structure in Attention.” Nature Reviews Neuroscience 19 (1): 34–48.

Noda, Takahiro, Tomoki Amemiya, Tomoyo I. Shiramatsu, and Hirokazu Takahashi. 2017. “Stimulus Phase Locking of Cortical Oscillations for Rhythmic Tone Sequences in Rats.” Frontiers in Neural Circuits 11 (January): 2.

Pandya, D. N., D. L. Rosene, and A. M. Doolittle. 1994. “Corticothalamic Connections of Auditory-Related Areas of the Temporal Lobe in the Rhesus Monkey.” The Journal of Comparative Neurology 345 (3): 447–71.

Parras, Gloria G., Javier Nieto-Diego, Guillermo V. Carbajal, Catalina Valdés-Baizabal, Carles Escera, and Manuel S. Malmierca. 2017. “Neurons along the Auditory Pathway Exhibit a Hierarchical Organization of Prediction Error.” Nature Communications 8 (1): 2148.

Paxinos, George, and Charles Watson. 2007. The Rat Brain in Stereotaxic Coordinates. Elsevier Science & Technology.

Polley, Daniel B., Heather L. Read, Douglas A. Storace, and Michael M. Merzenich. 2007. “Multiparametric Auditory Receptive Field Organization across Five Cortical Fields in the Albino Rat.” Journal of Neurophysiology 97 (5): 3621–38.

Rauschecker, J. P., and B. Tian. 2000. “Mechanisms and Streams for Processing of ‘What’ and ‘Where’ in Auditory Cortex.” Proceedings of the National Academy of Sciences of the United States of America 97 (22): 11800–11806.

Rosskothen-Kuhl, Nicole, Alexa N. Buck, Kongyan Li, and Jan Wh Schnupp. 2021. “Microsecond Interaural Time Difference Discrimination Restored by Cochlear Implants after Neonatal Deafness.” eLife 10 (e59300): e59300.

Schroeder, Charles E., and Peter Lakatos. 2009. “Low-Frequency Neuronal Oscillations as Instruments of Sensory Selection.” Trends in Neurosciences 32 (1): 9–18.

Szymanski, Francois D., Jose A. Garcia-Lazaro, and Jan W. H. Schnupp. 2009. “Current Source Density Profiles of Stimulus-Specific Adaptation in Rat Auditory Cortex.” Journal of Neurophysiology 102 (3): 1483– 90.

Szymanski, Francois D., Neil C. Rabinowitz, Cesare Magri, Stefano Panzeri, and Jan W. H. Schnupp. 2011. “The Laminar and Temporal Structure of Stimulus Information in the Phase of Field Potentials of Auditory Cortex.” The Journal of Neuroscience: The Official Journal of the Society for Neuroscience 31 (44): 15787–801.

Tao, Can, Guangwei Zhang, Chang Zhou, et al. 2017. “Diversity in Excitation-Inhibition Mismatch Underlies Local Functional Heterogeneity in the Rat Auditory Cortex.” Cell Reports 19 (3): 521–31.

Ten Oever, Sanne, and Andrea E. Martin. 2024. “Interdependence of ‘What’ and ‘When’ in the Brain.” Journal of Cognitive Neuroscience 36 (1): 167–86.

Ulanovsky, Nachum, Liora Las, Dina Farkas, and Israel Nelken. 2004. “Multiple Time Scales of Adaptation in Auditory Cortex Neurons.” The Journal of Neuroscience: The Official Journal of the Society for Neuroscience 24 (46): 10440–53.

Valt, Christian, Fran López-Caballero, Angelantonio Tavella, et al. 2024. “Abnormal Inter-Hemispheric Effective Connectivity from Left to Right Auditory Regions during Mismatch Negativity (MMN) Tasks in Psychosis.” Psychiatry Research 342 (116189): 116189.

De Venecia, R. K., C. B. Smelser, S. D. Lossman, and N. T. McMullen. 1995. “Complementary Expression of Parvalbumin and Calbindin D-28k Delineates Subdivisions of the Rabbit Medial Geniculate Body.” The Journal of Comparative Neurology 359 (4): 595–612.

Wacongne, Catherine, Jean-Pierre Changeux, and Stanislas Dehaene. 2012. “A Neuronal Model of Predictive Coding Accounting for the Mismatch Negativity.” The Journal of Neuroscience: The Official Journal of the Society for Neuroscience 32 (11): 3665–78.

Winer, Jeffery A., Lee M. Miller, Charles C. Lee, and Christoph E. Schreiner. 2005. “Auditory Thalamocortical Transformation: Structure and Function.” Trends in Neurosciences 28 (5): 255–63.

